# A Computer-aided Method for Identifying the Presence of Softwood Growth Ring Boundaries

**DOI:** 10.1101/2020.06.23.166702

**Authors:** Qi-Zhao Lin, Tuo He, Yong-Ke Sun, Xin He, Jian Qiu

## Abstract

The objective of this study was to develop a computer-aided method to quantify the obvious degree of growth ring boundaries of softwood species, based on data analysis with some image processing technologies. For this purpose, a 5× magnified cross-section color micro-image of softwood was cropped into 20 sub-images, then every image is binarized as a gray image according to an automatic threshold value. After that, the number of black pixels in the gray image was counted row by row and the number of black pixels was binarized to 0 or 100. Finally, a transition band from earlywood to latewood on the sub-image was identified. If this was successful, the growth ring boundaries of the sub-image are distinct, otherwise they were indistinct or absent. If 10 of the 20 sub-images are distinct, with the majority voting method, the growth ring boundaries of softwood are distinct, otherwise they are indistinct or absent. The proposed method has been visualized as a growth-ring-boundary detecting system based on the .NET Framework. A sample of 100 micro-images (Supplementary Images) of softwood cross-sections were selected for evaluation purposes. In short, this detecting system computes the obvious degree of growth ring boundaries of softwood species by image processing involved image importing, image cropping, image reading, image grayscale, image binarization, data analysis. The results showed that the method used avoided mistakes made by the manual comparison method of identifying the presence of growth ring boundaries, and it has a high accuracy of 98%.

## INTRODUCTION

Generally, a wood species can be identified according to the macroscopic and microscopic structural characteristics of the wood, which is a time-consuming process. The traditional methods of wood identification include manual comparison, dichotomous keys, multiple entry keys, punch card search and computer database program search (Wheeler & Baas 1998). Researchers have also tried to use DNA molecular marker technology (Yu *et al*. 2017), near-infrared spectroscopy technology (Snel *et al*. 2018), GC-MS technology (Wang *et al*. 2018), computer vision technology (Hwang *et al*. 2018) and other auxiliary identification methods to improve the accuracy of traditional methods in wood identification and speed the process of identifying wood species.

Recently, researchers have attempted to recognize wood species by utilizing a growth ring boundary detection algorithm (Fahrurozi *et al*. 2016) such as the Gray Level Co-occurrence Matrix (Xie & Wang 2015; Fahrurozi *et al*. 2016), and the color histogram statistical method (Zhao 2013) to extract wood features. Subsequently, various techniques, including Support Vector Machine (SVM) (Sun *et al*. 2015), K-nearest neighbor (KNN) (Gani & Mohamed 2013; Kobayashi *et al*. 2015; Fuentealba *et al*. 2005), and neural network (Zhao *et al*. 2014; Yuce *et al*. 2014), have been used to create many classifiers.

According to the *IAWA list of microscopic features for softwood identification* (IAWA Committee. 2004. IAWA list of microscopic features for softwood identification. IAWA Journal, 25(1): 1-70.), *Tsuga chinensis* var*. forrestii* (Fig.1) is always identified as having distinct growth ring boundaries, but *Podocarpus neriifolius* (Fig.2) may be recognized as having either obvious growth ring boundaries or not obvious growth ring boundaries (Jiang *et al*. 2010). The presence of growth ring boundaries in *Podocarpus neriifolius* varies from person to person, due to definitions of “*growth ring boundaries=growth rings with an abrupt structural change at boundaries between them*” and “*growth ring boundaries indistinct or absent=growth rings boundaries vague and with marked gradual structural changes*” being qualitative, not quantitative, which generates a serious problem for a wood identification researcher.

**Figure 1.**
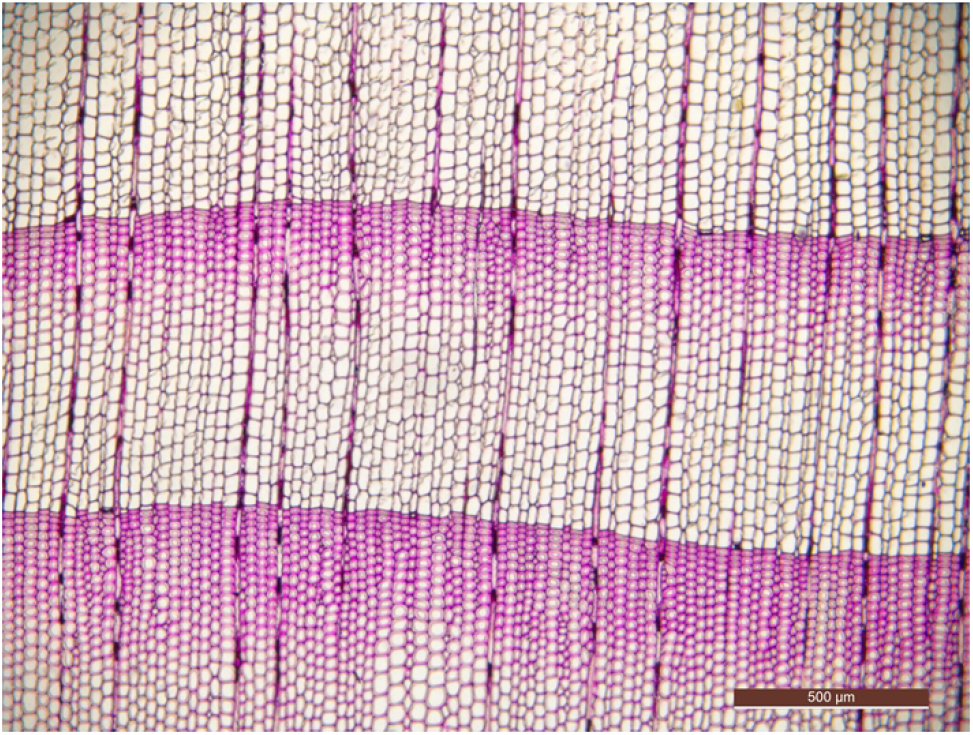
Cross-section of *Tsuga chinensis* var. *forrestii*.

**Figure 2.**
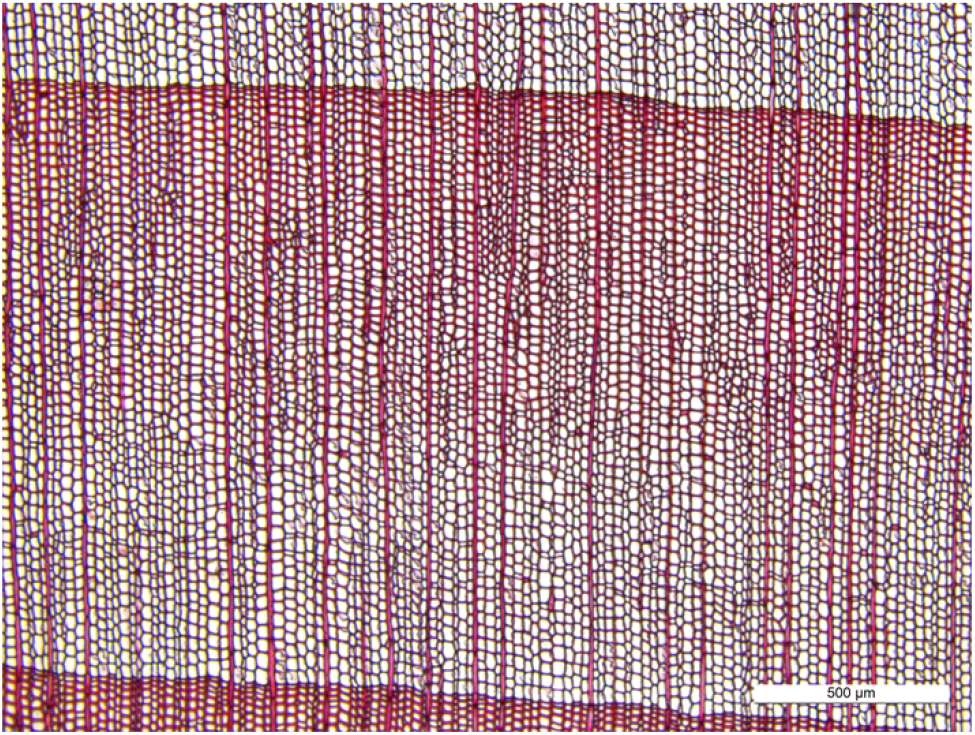
Cross-section of *Podocarpus neriifolius*.

There are no tools available to quantify the obvious degree of growth ring boundaries in softwood species, although many tools can be used to analyze wood anatomy images, for example, the image analysis tool ROXAS can be used to recognize annual rings in large samples (linear and circular) with > 100 annual rings (see www.wsl.ch/roxas), and to build centuries-long tracheid-lumen chronologies in conifers (Georg & Marco, 2013) or quantify plasticity in vessel grouping (Georg 2014). DENDRO-2003 densitometer can be used to measure the density profiles of tree ring (Vaganov *et al*. 2009). Besides, WinDENDRO can measure ring-width manually on sampled cross-sections (Wagner *et al*. 2010). They can not judge whether the softwood growth ring boundaries are obvious.

To address this issue, in this paper, it was proposed to develop a computer-aided method to identify quantitatively whether there are distinct growth ring boundaries present in softwood species, and provide a powerful quantitative wood anatomy tool (Georg V A. *et al*. 2016), in contrast to the method of identifying tree species (Yuce *et al*. 2014; Fahrurozi *et al*. 2016; Fahrurozi *et al*. 2016; Fuentealba *et al*. 2005; Gani & Mohamed 2013; Kobayashi *et al*. 2015; Sun *et al*. 2015; Xie & Wang. 2015; Zhao 2013; Zhao *et al*. 2014).

## MATERIALS AND METHODS

### Image acquisition

A total of 100 microscopic slides were collected from Wood Collections, Chinese Academy of Forestry, representing 100 species (Supplementary Table 1), involving 8 families of Ginkgoaceae, Araucariaceae, Podocarpaceae, Cephalotaxaceae, Taxaceae, Pinaceae, Taxodiaceae, and Cupressaceae. Imaging was performed with a digital camera (LEICA DMC4500) mounted on a light microscope (LEICA DM2000 LED). Images of 2560 × 1920 pixels were captured at 5× magnification using Leica Application Suite (Version 4.9.0). Numerical analysis and data visualization were carried out using Origin8.0.

### Proposed methods

The workflow of this study is presented in Fig. 3. The description of the method includes three parts. 1) The flow chart of the method 2) The image processing techniques used in this study 3) How to obtain the final result by some digital techniques, which also include data and statistical analysis. A brief introduction of the workflow is given below. Based on the workflow, a visual computer program has been designed by authors.

**Figure 3.**
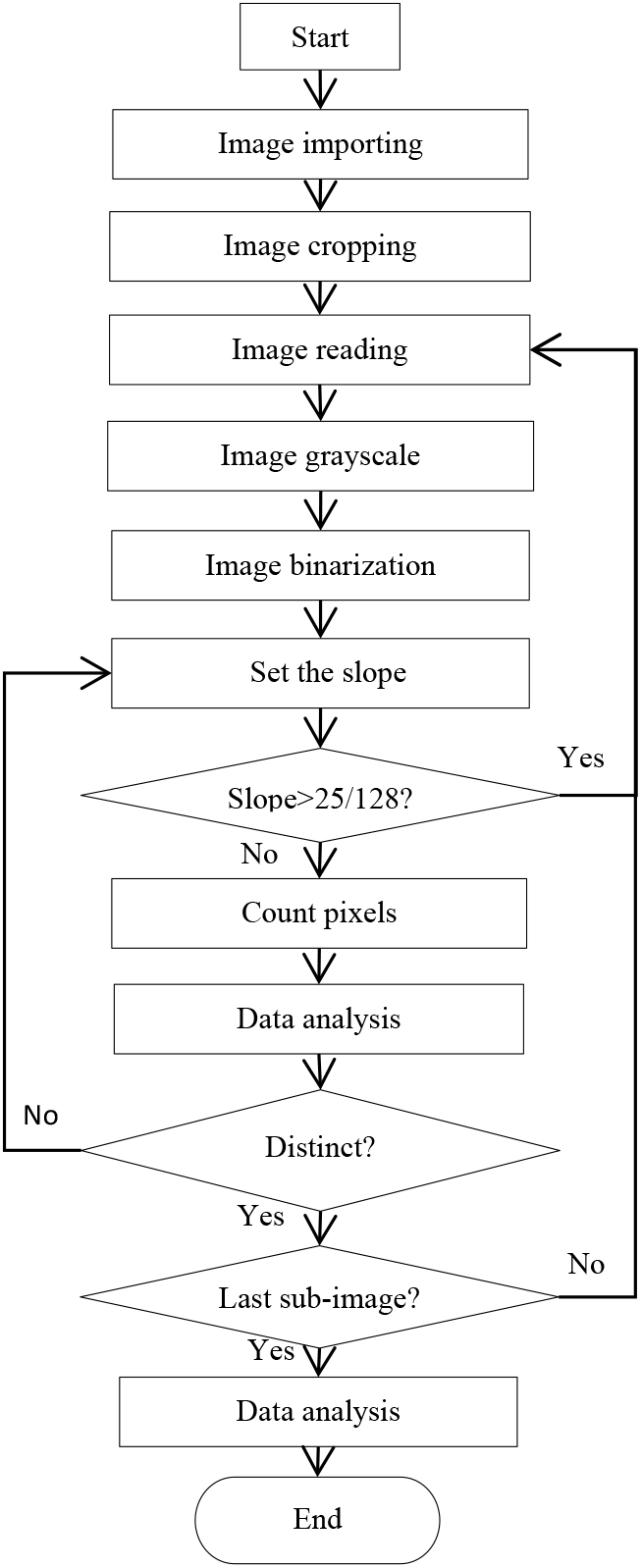
Flow chart of the program.

## PART I: FLOW CHART

The flow chart was described below:

Step 1.Input a microscopy RGB (RGB, R=Red, G=Green, B=Blue) color image collected from cross-section of softwood. The growth rings are parallel to the horizontal direction as far as possible.
Step 2.Crop imported image into 20 sub-images averagely in size along the horizontal direction.
Step 3.Read the images in sequence. If it is successful, then turn to Step 4, otherwise switch to Step 9.
Step 4.Convert a color image into a grayscale image.
Step 5.Calculate a threshold, and then change the grayscale image to get a binary image.
Step 6.Set up the slope value in a loop, if the slope is bigger than the threshold value, turn to Step 3.
Step 7.Count the number of black pixels in each row of the binary image from top to bottom by the slope.
Step 8.Analyze the data generated from Step 7, and then find a row index that meets the specific criteria described below. If it can be found, the growth ring boundaries are distinct and turn to Step 3, otherwise, the growth ring boundaries are indistinct or absent, and turn to Step 6.
Step 9.Statistically analyze all the results output by Step 8. If at least 10 sub-images resulted in distinct growth ring boundaries, then the growth ring boundaries of the sample are distinct, otherwise, they are indistinct or absent.

To operate the program correctly, the detailed instructions and constraints are emphasized as follows.

1. At Step 1, a suitable image shown as Fig. 4 (A) is acceptable but the image like Fig.4 (B) cannot be used, since the slope of the growth ring boundary in Fig. 4 (B) is too large. The growth ring on input images should be horizontal. To avoid finding wood rays as the boundary of the growth ring, the maximum acceptable slope of the growth ring boundary designed by the proposed computer-aided method is 0.195 (25 / 128).
2. At Step 2, compared with the original imported image, the sub-image after cropping can reduce the ordinate range of the boundary of the growth ring. Fig. 5 (A) shows an original image of a 5× magnified microscopy image of the cross-section of *Taxus wallichiana* without a scale, and the Fig.5 (B) shows 20 sub-images after image cropping.

**Figure 4.**
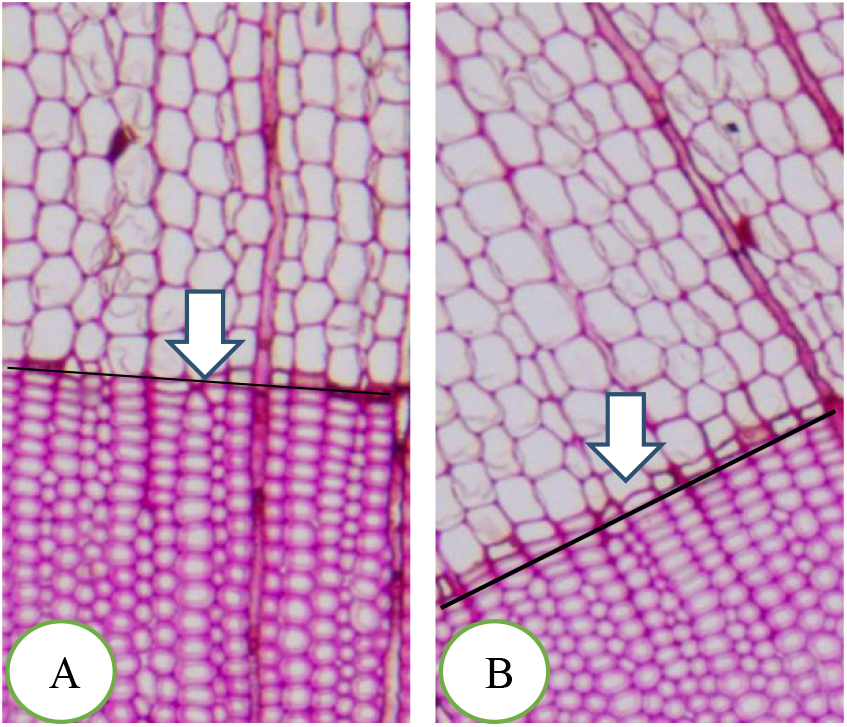
Examples of imported images of local regions: -A: Suitable image. - B: Unsuitable image.

**Figure 5.**
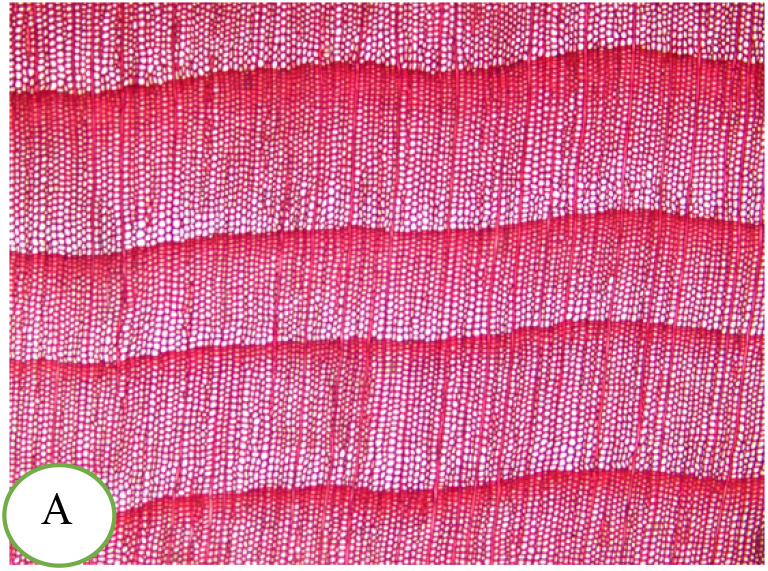
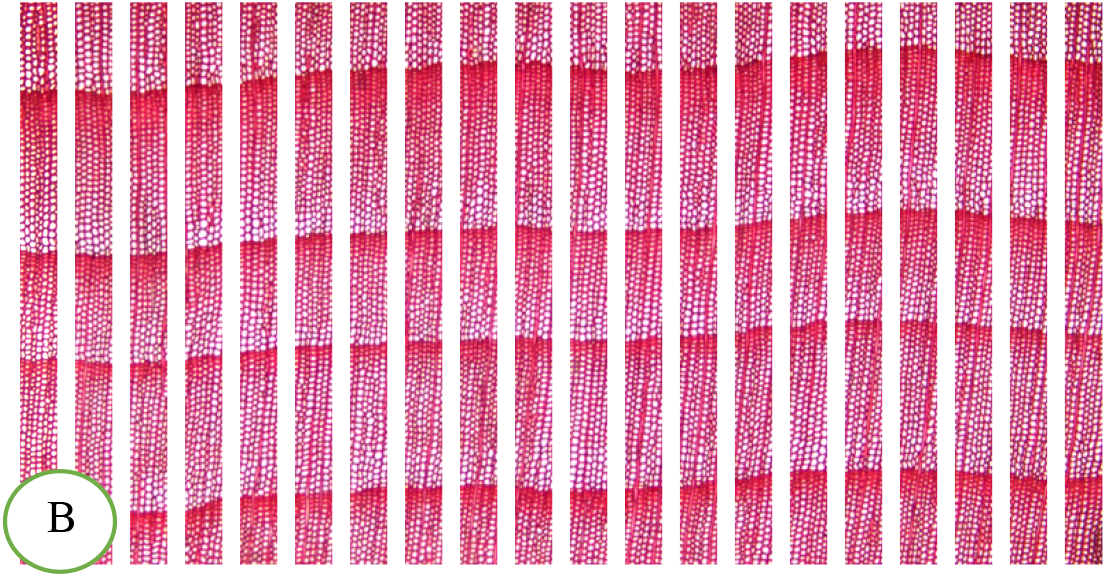
Example of image division: -A: Original image. -B: 20 sub-image set.

## PART II: IMAGE PROCESSING TECHNOLOGY

At Step 4, the image processing is performed to find growth rings with an abrupt change at the boundaries. The microscopic RGB images of a cross-section of a softwood species were first stored in a two-dimensional matrix defined as *f(x,y)*, converted into a grayscale image, and then changed into a black-and-white binary image. Every pixel point of the color image was calculated by Equation 1 below. The gray value of R, G, B ranges from 0 to 255, and values of all these pixel points lay between 0 to 255 calculated by Equation 1. Image’s dark pixel points represent the tracheid wall thickness, due to tracheid wall is dark but tracheid lumen is light.

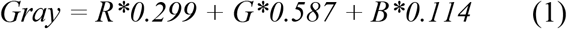

For segmenting an image, setting up the threshold was simple, efficient, and fast (Mehmet & Bülent 2004). At Step 5, thresholds were calculated by the program designed by the authors. Threshold values may be changed with different grayscale images. For getting the threshold τ, the program calculated the average value μ and standard deviation σ of all pixel points. These parameters μ, σ, and τ were calculated by Equations 2, 3, and 4, respectively.

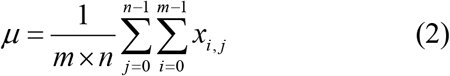

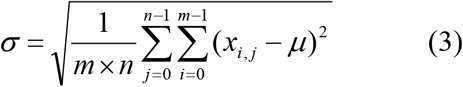

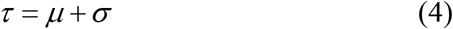

Where *x*_*i,j*_ is the value of a grayscale image pixel point;

*i* is the row index
*j* is the column index;
*m* is the image height;
*n* is the image width.

The program output various thresholds from different binary images, as shown in Fig. 5. By Equation 5, the value of pixel *p(x)* is defined as 0 (black) when if the gray value is less than the threshold τ, otherwise it is defined as 255 (white). After this process, a grayscale image could be converted into a binary image finally.

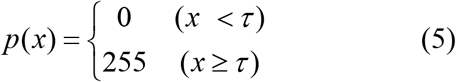

## PART III: MATHEMATICAL TECHNIQUE

A mathematical technique is conducted at Steps 7-9. Fig. 6 shows an example of counting black pixels. The first column of the sheet contains row index and the second column contains the counting of black pixels. The first row is at the top of the binary image. The number of black pixels in each row *Y*_*j*_ is counted by Equation 6, To find the growth ring boundary, *j* was corrected by Equation 7, where *s* was the slope of the growth ring boundary and computed by Equation 8. The proposed method gets *k* in a loop process. The maximum *j* equaled to *h*-*w*, where *h* is the height of the image.

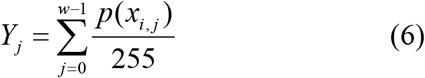

Where: *i* is the row index; *j* is the column index; and *w* is the width of the image.

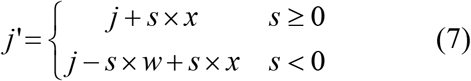

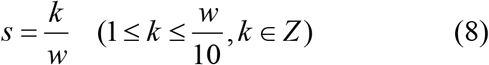

**Figure 6.**
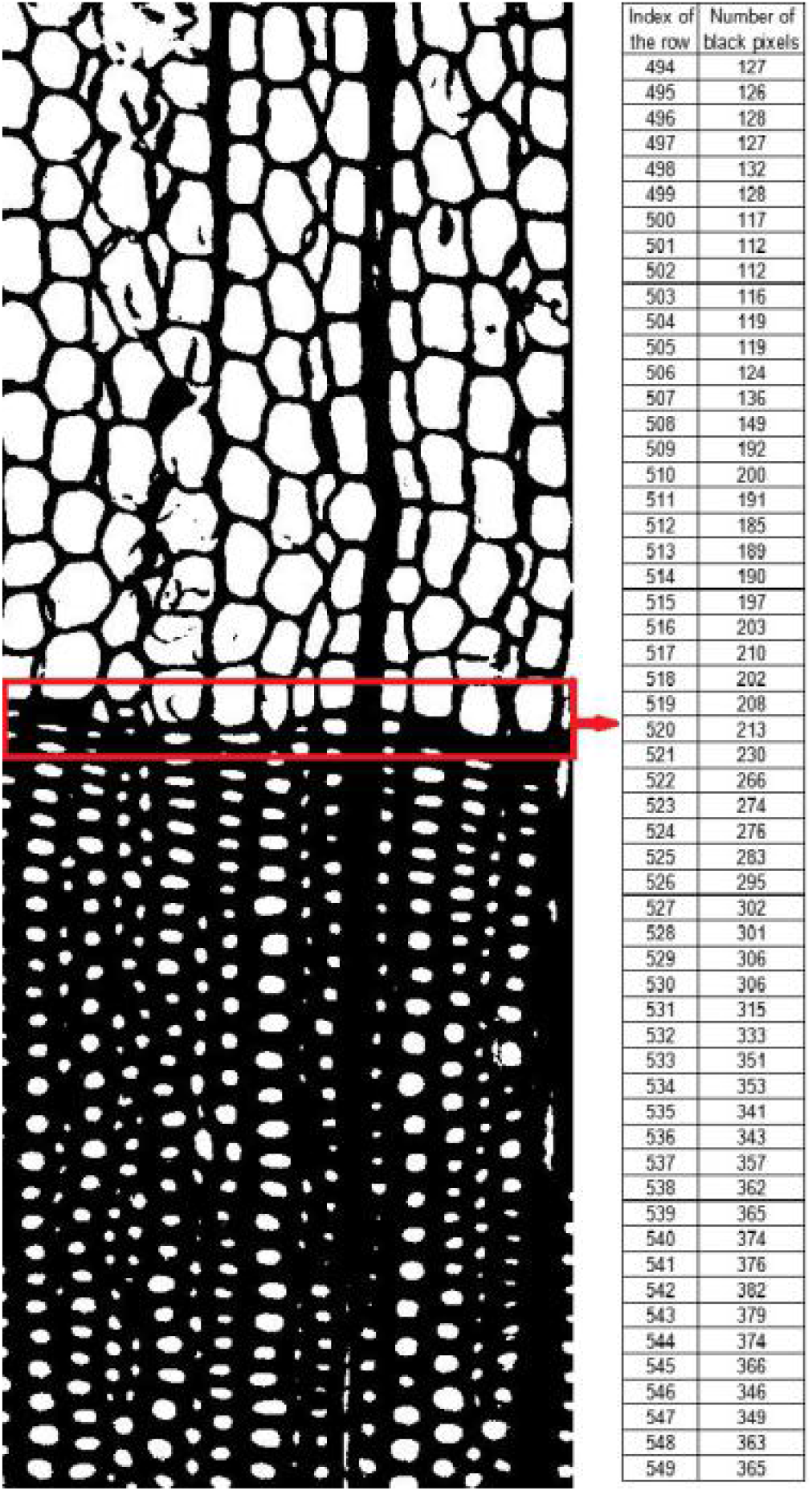
Example of counting black pixels.

After Step 7, the computer program normalizes these values by Equation 9 at Step 8.

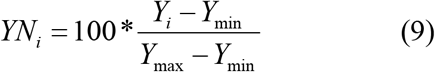

Where: *YN*_*i*_ is the normalized value of *Y*_*i*_;

*i, is* the row index
*Y*_*min*_ *is* the minimum value of *y*
*Y*_*max*_ is the maximum value of *y*.

Fig. 8 and Fig. 9 differ in the longitudinal coordinates. The purpose of normalization operation is to make all scatter plots have the same longitudinal coordinates ranging that from 0 to 100.

**Figure 8.**
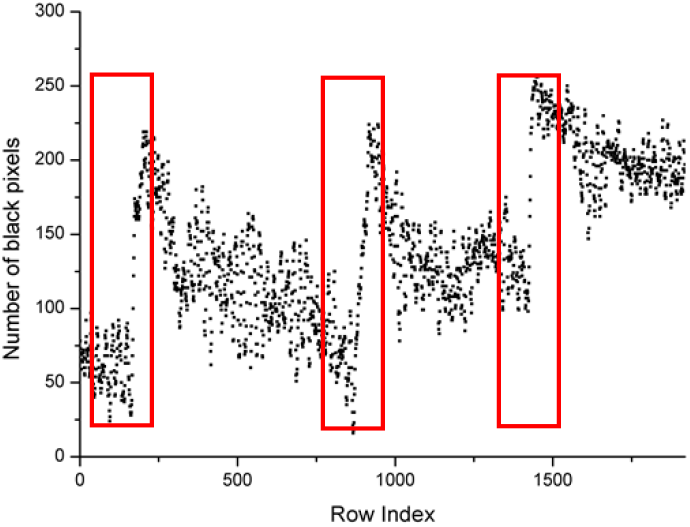
Scatter plot of the number of black pixels of Fig. 7 (B)

**Figure 9.**
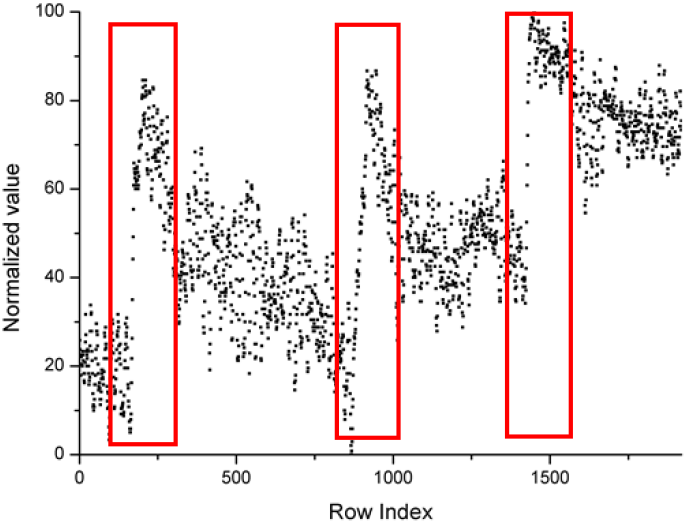
Scatter plot of the number of black pixels of Fig. 7 (B) after normalization.

The computer program calculates normalized values by Equation 10 and processes normalized values to 0 or 100, which is a binarized operation.

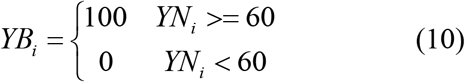

Where *YB*_*i*_ is the binarized value of *YN*_*i*_ *i* is the row index.

Fig. 10 is a scatter plot of the number of black pixels in Fig. 7 (B) after being binarized. From Fig. 8 to Fig. 10, it was easy to find out special regions that are labeled by the red rectangular box. These special regions represent growth ring boundaries.

**Figure 7.**
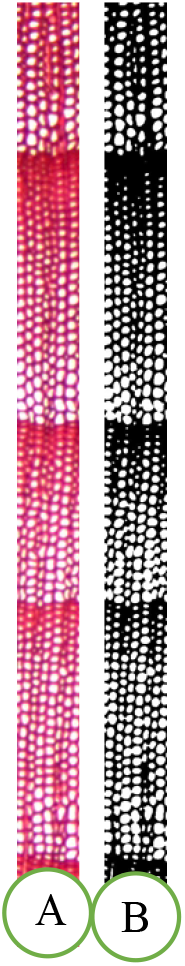
An example image of *Taxus wallichiana*.

**Figure 10.**
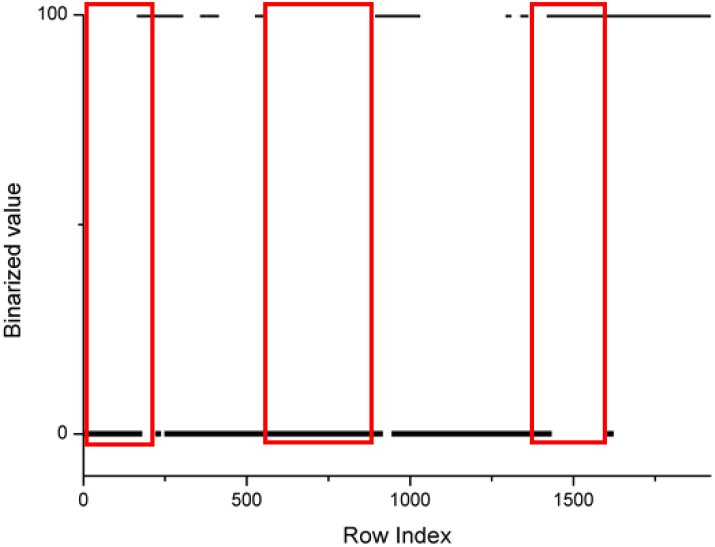
Scatter plot of the number of black pixels of Fig. 7 (B) after binarization.

In this study, a sub-method is included at Step 8, which aims to find a row index making *YB*_*i*_ = 0, *YB*_*i+*_ = 100, 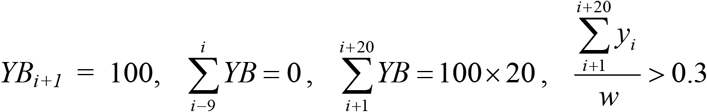 and 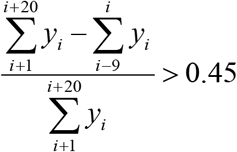. If it can be found, the growth ring boundaries are defined as “distinct”, otherwise, they are indistinct or absent.

At Step 9, the computer program analyzed the results generated from Step 8. If at least 10 sub-images are reported as “distinct”, then the growth ring boundaries of the sample were distinct, otherwise, they were indistinct or absent.

## RESULTS & DISCUSSION

For using this method, a visual computer program as shown in Fig. 11 was designed with C# based on the .NET Framework. The experimental results are shown in Table 1 including 100 softwood species. All the cross-section micro-images (Supplementary Images) are 2560×1920 pixels with the same magnification.

**Figure 11.**
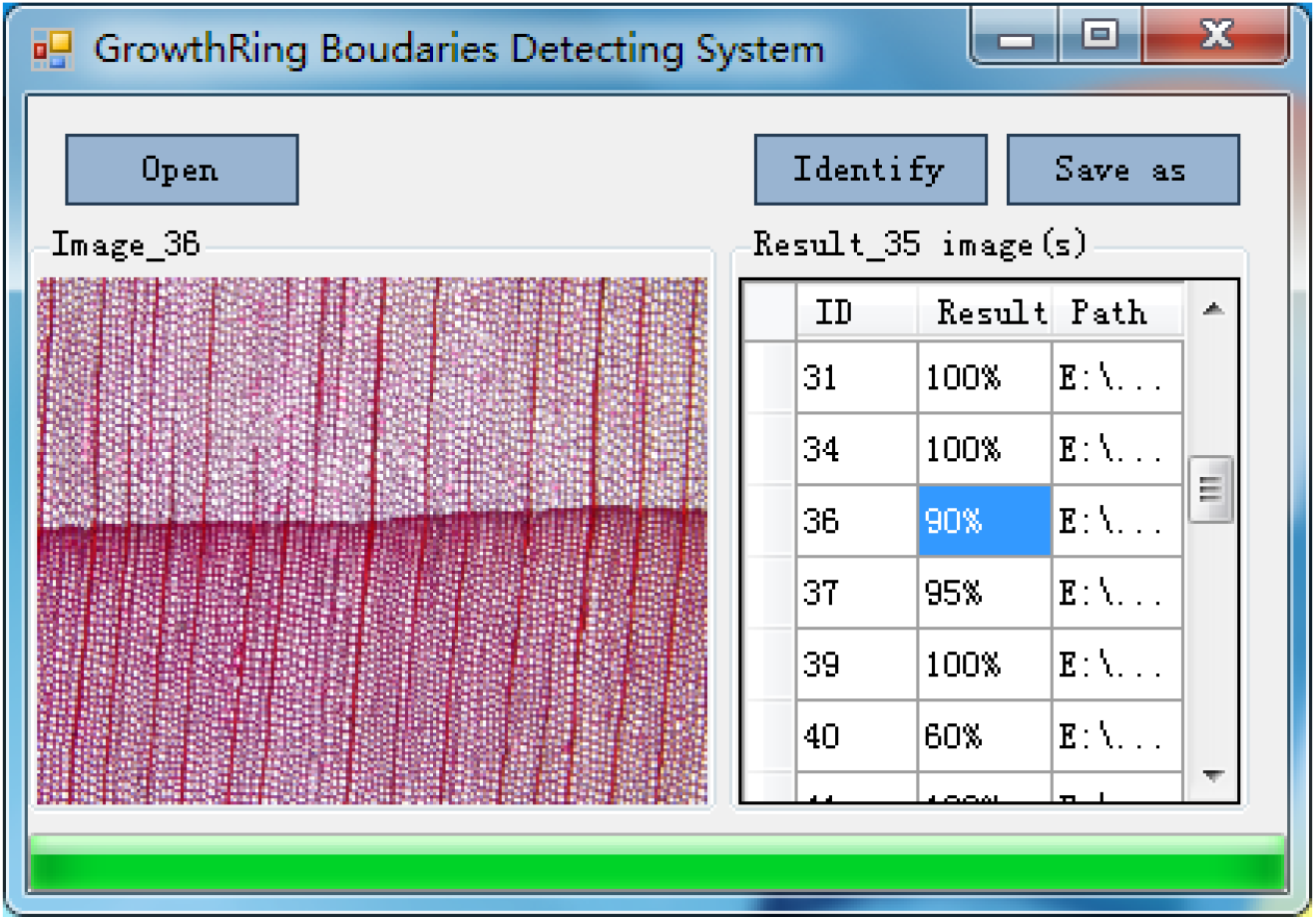
The interface of the program.

**Table 1.**
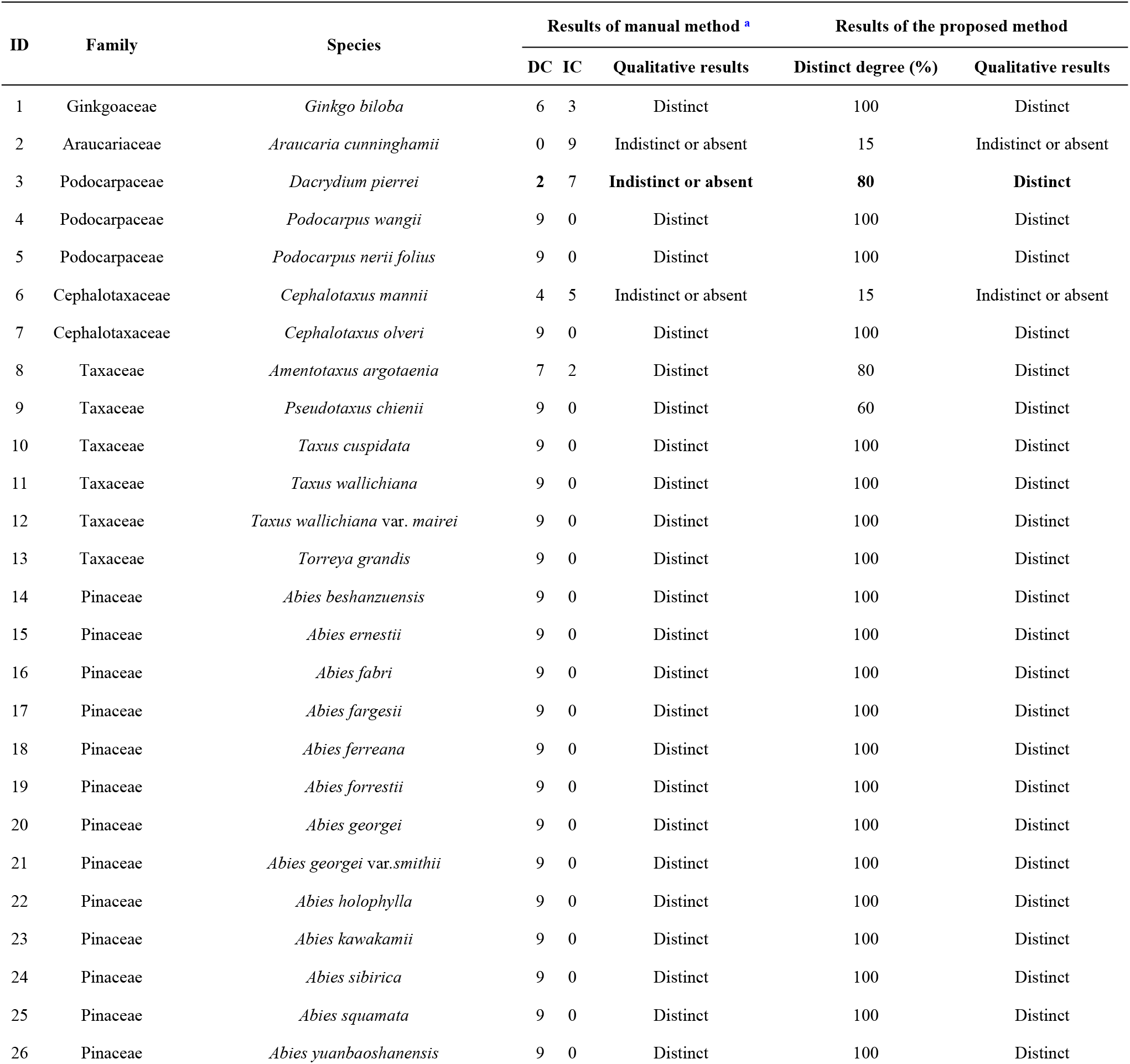
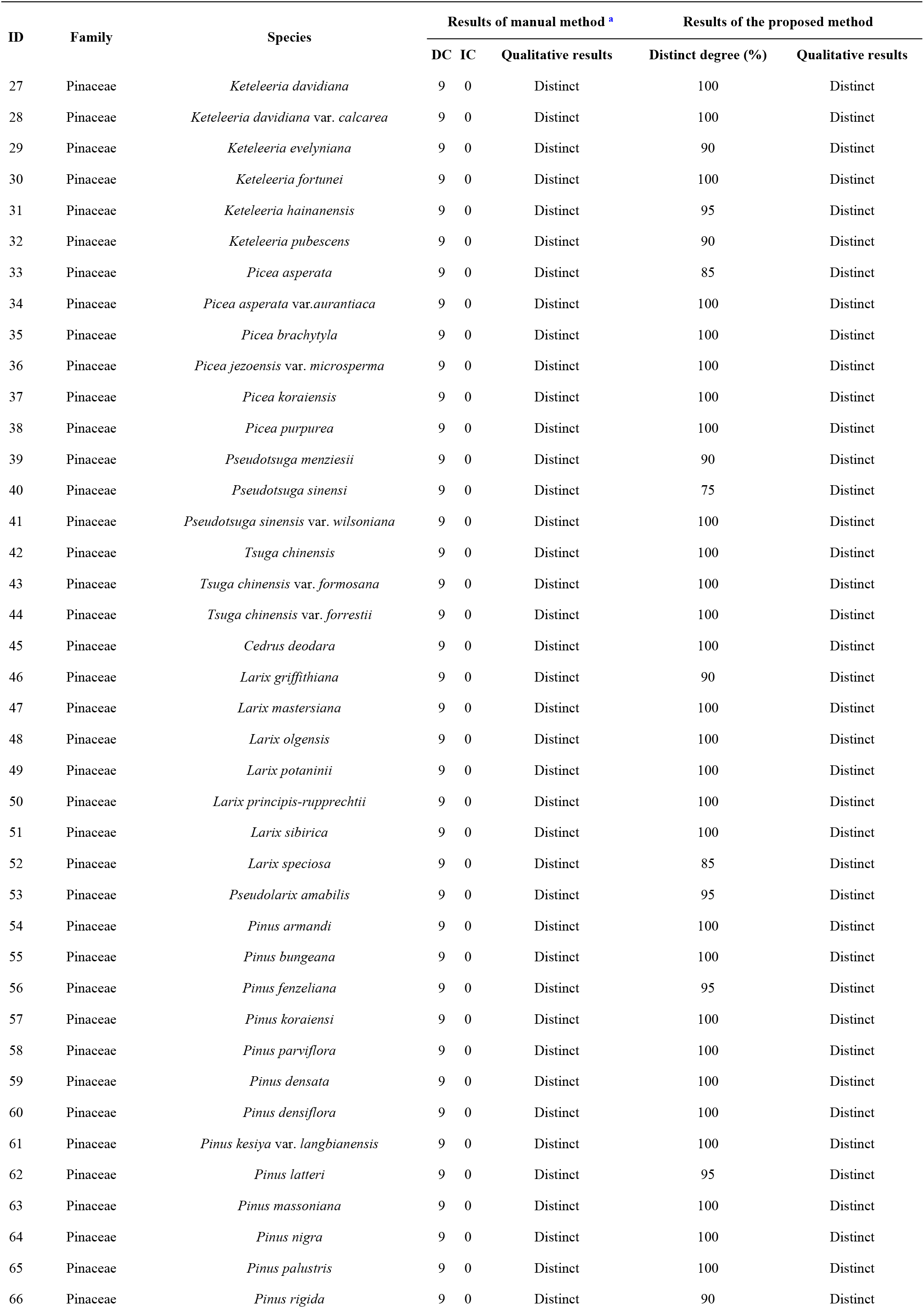
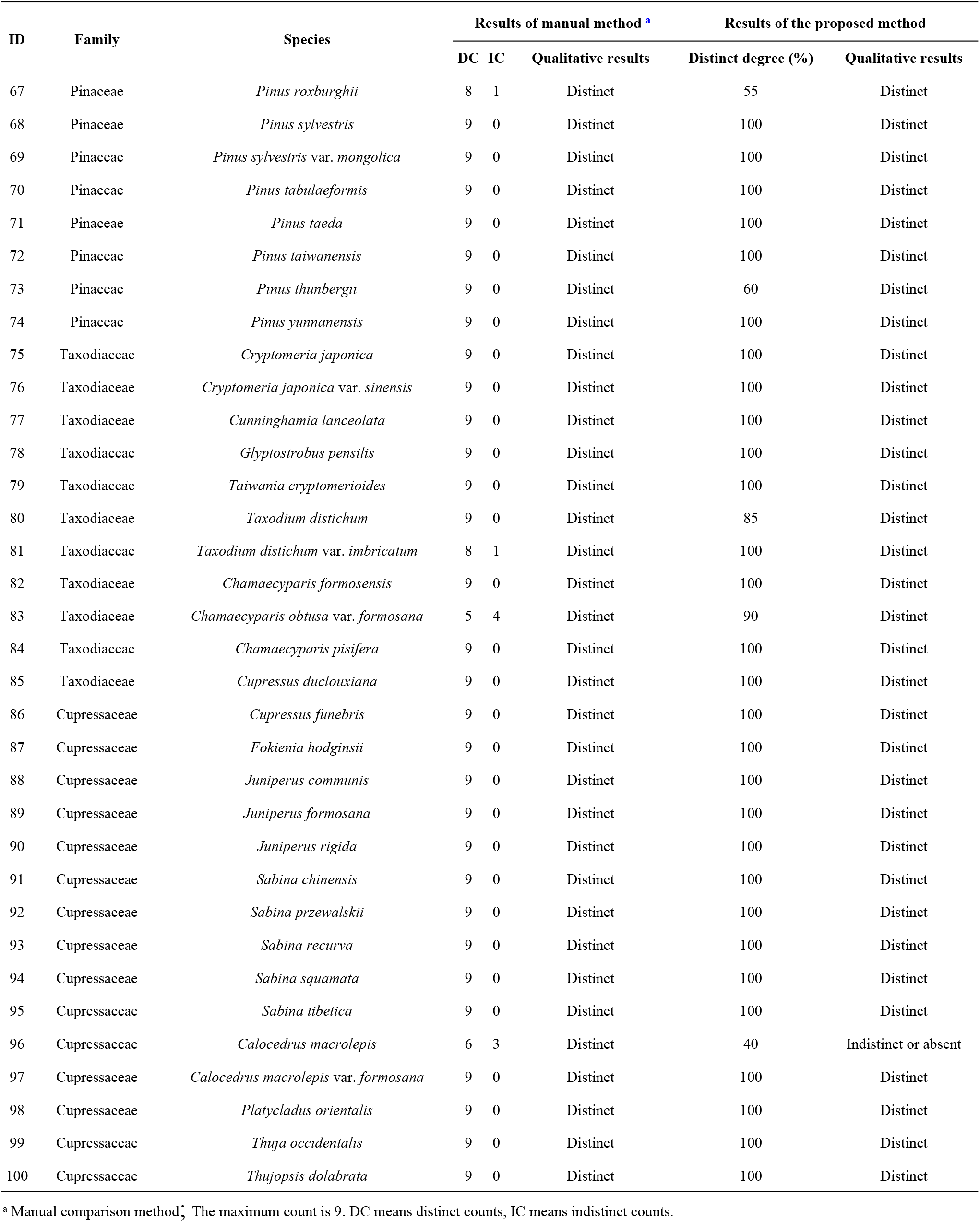
The experimental results of 100 softwood species

| ID | Family | Species | Results of manual method <sup>a</sup> |  |  | Results of the proposed method |  |
| --- | --- | --- | --- | --- | --- | --- | --- |
|  |  |  | DC | IC | Qualitative results | Distinct degree (%) | Qualitative results |
| 1 | Ginkgoaceae | *Ginkgo biloba* | 6 | 3 | Distinct | 100 | Distinct |
| 2 | Araucariaceae | *Araucaria cunninghamii* | 0 | 9 | Indistinct or absent | 15 | Indistinct or absent |
| 3 | Podocarpaceae | *Dacrydium pierrei* | 2 | 7 | **Indistinct or absent** | **80** | **Distinct** |
| 4 | Podocarpaceae | *Podocarpus wangii* | 9 | 0 | Distinct | 100 | Distinct |
| 5 | Podocarpaceae | *Podocarpus nerii folius* | 9 | 0 | Distinct | 100 | Distinct |
| 6 | Cephalotaxaceae | *Cephalotaxus mannii* | 4 | 5 | Indistinct or absent | 15 | Indistinct or absent |
| 7 | Cephalotaxaceae | *Cephalotaxus olveri* | 9 | 0 | Distinct | 100 | Distinct |
| 8 | Taxaceae | *Amentotaxus argotaenia* | 7 | 2 | Distinct | 80 | Distinct |
| 9 | Taxaceae | *Pseudotaxus chienii* | 9 | 0 | Distinct | 60 | Distinct |
| 10 | Taxaceae | *Taxus cuspidata* | 9 | 0 | Distinct | 100 | Distinct |
| 11 | Taxaceae | *Taxus wallichiana* | 9 | 0 | Distinct | 100 | Distinct |
| 12 | Taxaceae | *Taxus wallichiana* var. *mairei* | 9 | 0 | Distinct | 100 | Distinct |
| 13 | Taxaceae | *Torreya grandis* | 9 | 0 | Distinct | 100 | Distinct |
| 14 | Pinaceae | *Abies beshanzuensis* | 9 | 0 | Distinct | 100 | Distinct |
| 15 | Pinaceae | *Abies ernestii* | 9 | 0 | Distinct | 100 | Distinct |
| 16 | Pinaceae | *Abies fabri* | 9 | 0 | Distinct | 100 | Distinct |
| 17 | Pinaceae | *Abies fargesii* | 9 | 0 | Distinct | 100 | Distinct |
| 18 | Pinaceae | *Abies ferreana* | 9 | 0 | Distinct | 100 | Distinct |
| 19 | Pinaceae | *Abies forrestii* | 9 | 0 | Distinct | 100 | Distinct |
| 20 | Pinaceae | *Abies georgei* | 9 | 0 | Distinct | 100 | Distinct |
| 21 | Pinaceae | *Abies georgei* var. *smithii* | 9 | 0 | Distinct | 100 | Distinct |
| 22 | Pinaceae | *Abies holophylla* | 9 | 0 | Distinct | 100 | Distinct |
| 23 | Pinaceae | *Abies kawakamii* | 9 | 0 | Distinct | 100 | Distinct |
| 24 | Pinaceae | *Abies sibirica* | 9 | 0 | Distinct | 100 | Distinct |
| 25 | Pinaceae | *Abies squamata* | 9 | 0 | Distinct | 100 | Distinct |
| 26 | Pinaceae | *Abies yuanbaoshanensis* | 9 | 0 | Distinct | 100 | Distinct |
|  |  |  | DC | IC | Qualitative results | Distinct degree (%) | Qualitative results |
| 27 | Pinaceae | *Keteleeria davidiana* | 9 | 0 | Distinct | 100 | Distinct |
| 28 | Pinaceae | *Keteleeria davidiana* var. *calcarea* | 9 | 0 | Distinct | 100 | Distinct |
| 29 | Pinaceae | *Keteleeria evelyniana* | 9 | 0 | Distinct | 90 | Distinct |
| 30 | Pinaceae | *Keteleeria fortunei* | 9 | 0 | Distinct | 100 | Distinct |
| 31 | Pinaceae | *Keteleeria hainanensis* | 9 | 0 | Distinct | 95 | Distinct |
| 32 | Pinaceae | *Keteleeria pubescens* | 9 | 0 | Distinct | 90 | Distinct |
| 33 | Pinaceae | *Picea asperata* | 9 | 0 | Distinct | 85 | Distinct |
| 34 | Pinaceae | *Picea asperata* var. *aurantiaca* | 9 | 0 | Distinct | 100 | Distinct |
| 35 | Pinaceae | *Picea brachytyla* | 9 | 0 | Distinct | 100 | Distinct |
| 36 | Pinaceae | *Picea jezoensis* var. *microsperma* | 9 | 0 | Distinct | 100 | Distinct |
| 37 | Pinaceae | *Picea koraiensis* | 9 | 0 | Distinct | 100 | Distinct |
| 38 | Pinaceae | *Picea purpurea* | 9 | 0 | Distinct | 100 | Distinct |
| 39 | Pinaceae | *Pseudotsuga menziesii* | 9 | 0 | Distinct | 90 | Distinct |
| 40 | Pinaceae | *Pseudotsuga sinensi* | 9 | 0 | Distinct | 75 | Distinct |
| 41 | Pinaceae | *Pseudotsuga sinensis* var. *wilsoniana* | 9 | 0 | Distinct | 100 | Distinct |
| 42 | Pinaceae | *Tsuga chinensis* | 9 | 0 | Distinct | 100 | Distinct |
| 43 | Pinaceae | *Tsuga chinensis* var. *formosana* | 9 | 0 | Distinct | 100 | Distinct |
| 44 | Pinaceae | *Tsuga chinensis* var. *forrestii* | 9 | 0 | Distinct | 100 | Distinct |
| 45 | Pinaceae | *Cedrus deodara* | 9 | 0 | Distinct | 100 | Distinct |
| 46 | Pinaceae | *Larix griffithiana* | 9 | 0 | Distinct | 90 | Distinct |
| 47 | Pinaceae | *Larix mastersiana* | 9 | 0 | Distinct | 100 | Distinct |
| 48 | Pinaceae | *Larix olgensis* | 9 | 0 | Distinct | 100 | Distinct |
| 49 | Pinaceae | *Larix potaninii* | 9 | 0 | Distinct | 100 | Distinct |
| 50 | Pinaceae | *Larix principis-rupprechtii* | 9 | 0 | Distinct | 100 | Distinct |
| 51 | Pinaceae | *Larix sibirica* | 9 | 0 | Distinct | 100 | Distinct |
| 52 | Pinaceae | *Larix speciosa* | 9 | 0 | Distinct | 85 | Distinct |
| 53 | Pinaceae | *Pseudolarix amabilis* | 9 | 0 | Distinct | 95 | Distinct |
| 54 | Pinaceae | *Pinus armandi* | 9 | 0 | Distinct | 100 | Distinct |
| 55 | Pinaceae | *Pinus bungeana* | 9 | 0 | Distinct | 100 | Distinct |
| 56 | Pinaceae | *Pinus fenzeliana* | 9 | 0 | Distinct | 95 | Distinct |
| 57 | Pinaceae | *Pinus koraiensi* | 9 | 0 | Distinct | 100 | Distinct |
| 58 | Pinaceae | *Pinus parviflora* | 9 | 0 | Distinct | 100 | Distinct |
| 59 | Pinaceae | *Pinus densata* | 9 | 0 | Distinct | 100 | Distinct |
| 60 | Pinaceae | *Pinus densiflora* | 9 | 0 | Distinct | 100 | Distinct |
| 61 | Pinaceae | *Pinus kesiya* var. *langbianensis* | 9 | 0 | Distinct | 100 | Distinct |
| 62 | Pinaceae | *Pinus latteri* | 9 | 0 | Distinct | 95 | Distinct |
| 63 | Pinaceae | *Pinus massoniana* | 9 | 0 | Distinct | 100 | Distinct |
| 64 | Pinaceae | *Pinus nigra* | 9 | 0 | Distinct | 100 | Distinct |
| 65 | Pinaceae | *Pinus palustris* | 9 | 0 | Distinct | 100 | Distinct |
| 66 | Pinaceae | *Pinus rigida* | 9 | 0 | Distinct | 90 | Distinct |
|  |  |  | DC | IC | Qualitative results | Distinct degree (%) | Qualitative results |
| 67 | Pinaceae | *Pinus roxburghii* | 8 | 1 | Distinct | 55 | Distinct |
| 68 | Pinaceae | *Pinus sylvestris* | 9 | 0 | Distinct | 100 | Distinct |
| 69 | Pinaceae | *Pinus sylvestris* var. *mongolica* | 9 | 0 | Distinct | 100 | Distinct |
| 70 | Pinaceae | *Pinus tabulaeformis* | 9 | 0 | Distinct | 100 | Distinct |
| 71 | Pinaceae | *Pinus taeda* | 9 | 0 | Distinct | 100 | Distinct |
| 72 | Pinaceae | *Pinus taiwanensis* | 9 | 0 | Distinct | 100 | Distinct |
| 73 | Pinaceae | *Pinus thunbergii* | 9 | 0 | Distinct | 60 | Distinct |
| 74 | Pinaceae | *Pinus yunnanensis* | 9 | 0 | Distinct | 100 | Distinct |
| 75 | Taxodiaceae | *Cryptomeria japonica* | 9 | 0 | Distinct | 100 | Distinct |
| 76 | Taxodiaceae | *Cryptomeria japonica* var. *sinensis* | 9 | 0 | Distinct | 100 | Distinct |
| 77 | Taxodiaceae | *Cunninghamia lanceolata* | 9 | 0 | Distinct | 100 | Distinct |
| 78 | Taxodiaceae | *Glyptostrobus pensilis* | 9 | 0 | Distinct | 100 | Distinct |
| 79 | Taxodiaceae | *Taiwania cryptomerioides* | 9 | 0 | Distinct | 100 | Distinct |
| 80 | Taxodiaceae | *Taxodium distichum* | 9 | 0 | Distinct | 85 | Distinct |
| 81 | Taxodiaceae | *Taxodium distichum* var. *imbricatum* | 8 | 1 | Distinct | 100 | Distinct |
| 82 | Taxodiaceae | *Chamaecyparis formosensis* | 9 | 0 | Distinct | 100 | Distinct |
| 83 | Taxodiaceae | *Chamaecyparis obtusa* var. *formosana* | 5 | 4 | Distinct | 90 | Distinct |
| 84 | Taxodiaceae | *Chamaecyparis pisifera* | 9 | 0 | Distinct | 100 | Distinct |
| 85 | Taxodiaceae | *Cupressus duclouxiana* | 9 | 0 | Distinct | 100 | Distinct |
| 86 | Cupressaceae | *Cupressus funebris* | 9 | 0 | Distinct | 100 | Distinct |
| 87 | Cupressaceae | *Fokienia hodginsii* | 9 | 0 | Distinct | 100 | Distinct |
| 88 | Cupressaceae | *Juniperus communis* | 9 | 0 | Distinct | 100 | Distinct |
| 89 | Cupressaceae | *Juniperus formosana* | 9 | 0 | Distinct | 100 | Distinct |
| 90 | Cupressaceae | *Juniperus rigida* | 9 | 0 | Distinct | 100 | Distinct |
| 91 | Cupressaceae | *Sabina chinensis* | 9 | 0 | Distinct | 100 | Distinct |
| 92 | Cupressaceae | *Sabina przewalskii* | 9 | 0 | Distinct | 100 | Distinct |
| 93 | Cupressaceae | *Sabina recurva* | 9 | 0 | Distinct | 100 | Distinct |
| 94 | Cupressaceae | *Sabina squamata* | 9 | 0 | Distinct | 100 | Distinct |
| 95 | Cupressaceae | *Sabina tibetica* | 9 | 0 | Distinct | 100 | Distinct |
| 96 | Cupressaceae | *Calocedrus macrolepis* | 6 | 3 | Distinct | 40 | Indistinct or absent |
| 97 | Cupressaceae | *Calocedrus macrolepis* var. *formosana* | 9 | 0 | Distinct | 100 | Distinct |
| 98 | Cupressaceae | *Platycladus orientalis* | 9 | 0 | Distinct | 100 | Distinct |
| 99 | Cupressaceae | *Thuja occidentalis* | 9 | 0 | Distinct | 100 | Distinct |
| 100 | Cupressaceae | *Thujopsis dolabrata* | 9 | 0 | Distinct | 100 | Distinct |

<sup>a</sup> Manual comparison method; The maximum count is 9. DC means distinct counts, IC means indistinct counts.

As shown in Table 1, the manual comparison method was composed of 9 experts with experience in the identification of softwood to determine whether the boundaries of softwood growth ring were distinct by cross-section micro-image. Among these 100 cross-sections of softwood identified by 9 experts, 91 were identified as an obvious feature of growth ring boundaries by all experts, 1 was identified as a non-obvious feature of growth ring boundaries by all experts, and the other 8 shown in Table 2 were not.

**Table 2.** Different judgments on the other 8 cross-sections

| ID | Family | Species | Results of the manual method |  |  | Results of the proposed method |  |
| --- | --- | --- | --- | --- | --- | --- | --- |
|  |  |  | DC | IC | Qualitative results | Distinct degree (%) | Qualitative results |
| 1 | Ginkgoaceae | *Ginkgo biloba* | 6 | 3 | Distinct | 100 | Distinct |
| 3 | Podocarpaceae | *Dacrydium pierrei* | 2 | 7 | Indistinct or absent | 80 | Distinct |
| 6 | Cephalotaxaceae | *Cephalotaxus mannii* | 4 | 5 | Indistinct or absent | 15 | Indistinct or absent |
| 8 | Taxaceae | *Amentotaxus argotaenia* | 7 | 2 | Distinct | 80 | Distinct |
| 67 | Pinaceae | *Pinus roxburghii* | 8 | 1 | Distinct | 55 | Distinct |
| 81 | Taxodiaceae | *Taxodium distichum* var. *imbricatum* | 8 | 1 | Distinct | 100 | Distinct |
| 83 | Taxodiaceae | *Chamaecyparis obtusa* var. *formosana* | 5 | 4 | Distinct | 90 | Distinct |
| 96 | Cupressaceae | *Calocedrus macrolepis* | 6 | 3 | Distinct | 40 | Indistinct or absent |

As shown in Table 2, different experts have different judgments on the transition type of the growth ring boundaries on these 8 cross-sections of softwood involving 7 families of Ginkgoaceae, Podocarpaceae, Cephalotaxaceae, Taxaceae, Pinaceae, Taxodiaceae, and Cupressaceae. The proposed method provides a quantitative value of the degree of distinctness of growth ring boundaries, then provides a qualitative conclusion with the majority voting method. Compared with the traditional method (IAWA Committee. 2004. IAWA list of microscopic features for softwood identification. IAWA Journal, 25(1): 1-70.), to judge whether the growth ring boundaries were distinct, the advantage of the proposed method is providing a qualitative conclusion with the majority voting method based on quantitative computation, which minimized mistakes made by the manual comparison method.

Compared with qualitative results of the manual comparison method which have been run by distinct counts and indistinct counts with the majority voting method, qualitative results of the proposed method were different on *Dacrydium pierrei* (N_O_. 3) of the Podocarpaceae family and *Calocedrus macrolepis* (N_O_. 96) of the Cupressaceae family after quantifying the distinctness of growth ring boundaries. In other words, the accuracy of the proposed method was 98% assuming that the results of manual comparison were all correct.

This method may be improved by combining with a variance which can be used to measure the fluctuation of the image region, to automatically identify the transition from earlywood to latewood in softwoods. It could also be applied to assess the distinctiveness of hardwood growth ring boundaries.

## CONCLUSIONS

The results show that the computer-aided method avoided making the same mistakes as manual identification of growth ring boundaries, with a high accuracy of 98%. Experiments show that the computer-aided method was robust.

## ACKNOWLEDGMENTS

This research is supported by the National Keypoint Research and Invention Program in 13th Five-Year, China, No. 2016YFD0600702. We are grateful to our colleagues, J.R. Gao, H.S. He, Y.G. Li, B. Luo, L. Qin, M.Y. Ran, X. Wang, Y.L. Wang and Y.S. Yang for their help in manually identifying the presence of 100 softwood growth ring boundaries. We especially thank Dr. Y.F. Yin and Dr. H. Wan, for their suggestions on the improvement of this paper.

## APPENDIX A. Supplementary data

Supplementary data to this article can be found online at https://github.com/senly2019/Lin-Qizhao/.

